# ESMFold Hallucinates Native-Like Protein Sequences

**DOI:** 10.1101/2023.05.23.541774

**Authors:** Jeliazko R. Jeliazkov, Diego del Alamo, Joel D. Karpiak

## Abstract

We describe attempts to design protein sequences by inverting the protein structure prediction algorithm ESMFold. State-of-the-art protein structure prediction methods achieve high accuracy by relying on evolutionary patterns derived from either multiple sequence alignments (AlphaFold, RosettaFold) or pretrained protein language models (PLMs; ESMFold, OmegaFold). In principle, by inverting these networks, protein sequences can be designed to fulfill one or more design objectives, such as high prediction confidence, predicted protein binding, or other geometric constraints that can be expressed with loss functions. In practice, sequences designed using an inverted AlphaFold model, termed AFDesign, contain unnatural sequence profiles shown to express poorly, whereas an inverted RosettaFold network has been shown to be sensitive to adversarial sequences. Here, we demonstrate that these limitations do not extend to neural networks that include PLMs, such as ESMFold. Using an inverted ESMFold model, termed ESM-Design, we generated sequences with profiles that are both more native-like and more likely to express than sequences generated using AFDesign, but less likely to express than sequences rescued by the structure-based design method ProteinMPNN. However, the safeguard offered by the PLM came with steep increases in memory consumption, preventing proteins greater than 150 residues from being modeled on a single GPU with 80GB VRAM. During this investigation, we also observed the role played by different sequence initialization schemes, with random sampling of discrete amino acids improving convergence and model quality over any continuous random initialization method. Finally, we showed how this approach can be used to introduce sequence and structure diversification in small proteins such as ubiquitin, while respecting the sequence conservation of active site residues. Our results highlight the effects of architectural differences between structure prediction networks on zero-shot protein design.

## 1 Introduction

The widespread adoption of AI-based tools over the past three years has defined the field of protein design [1]. In contrast with previous generations of protein sequence design methods, such as position-specific scoring matrices (PSSMs) [2] and Potts models [3], generative deep learning models such as protein language models (PLMs) eschew explicit probabilistic frameworks entirely [4]. Instead, PLMs directly learn the patterns underlying sequence variation in the training data, without explicit knowledge on the part of the user as to the precise nature of those patterns. A common training procedure involves masking stretches of a protein sequence that need to be guessed, with errors backpropagated across the model [5]. Despite the superficially trivial nature of this task, this self-supervised approach, when executed on training databases containing millions of naturally occurring protein sequences, can expose PLMs to sequence patterns that extend beyond those sought by the practitioner. For example, PLMs trained on large corpuses of protein sequence data demonstrate extraordinary zero-shot capabilities in tasks such as fitness prediction [6] and *de novo* protein design [7].

Several groups have successfully designed protein sequences using structural information by repurposing state-of-the-art protein structure prediction methods such as AlphaFold2 (AF2) and RosettaFold (RF) [8, 9, 10, 11]. The release of AF2 has revolutionized the field of protein structure prediction and spawned a rich ecosystem of competing and expansive methods. Sequence design is one such application that was not built into the method’s architecture. In contrast with the design strategies outlined above that operate exclusively on sequence without any explicit knowledge of structure, inverted AF2 and RF can optimize sequences by maximizing one of several predicted confidence metrics and/or geometric constraints using either Markov Chain Monte Carlo or backpropagation. Maximizing confidence in a predicted structure from a highly accurate method should lead to highly stable, well-folding sequences. In practice, however, sequences designed this way have been widely reported to express poorly, if at all [9, 12], indicating this is more akin to an adversarial attack on the network than an intended use case, which limits its adoption as an integrated method for structure-based protein design.

We hypothesize that one such workaround may be found in more recent structure prediction methods, which augment the architecture of AF2 and RF by prepending PLMs trained on large sequence databases [13, 14]. Unlike AF2 and RF, which determine sequence profiles using alignments provided at runtime, PLM-based structure prediction methods such as ESMFold and OmegaFold learn sequence patterns during training. This innovation allows them to predict the structure of proteins from a single sequence alone. We reason, however, that it may allow ESMFold to overcome the limitations, outlined above, that prevented adoption of AF2 and RF for structure-based sequence design of proteins.

Here, we describe the results of inverting the ESMFold neural network to enable protein design. By converting the integer representation of amino acids sequences to a continuous one-hot encoding, ESMFold was used to backpropagate structural losses into sequence modifications. We then evaluated several combinations of sequence initialization schemes, optimizers, and loss functions, and partially characterized their interplay. Strikingly, amino acid initialization played a key role in convergence and amino acid usage rates. The choice of loss function also affected convergence, and we demonstrate that backpropagation through ESMFold can minimize RMSD loss albeit taking 10-20 fold more steps to stably converge. We found that initializing with discrete amino acids by sampling a distribution similar to UniRef produces designs with the most “natural” sequences. The resulting sequences were both more native-like and more likely to express than sequences generated from an inverted AlphaFold network (AFDesign). However, ESM-Design sequences were judged less likely to express than sequences generated by the dedicated structure-based sequence design algorithm ProteinMPNN. Moreover, in cases where ESM-Design was used to generate diversity for a target enzyme, the designed sequences retained conserved active site residues without being explicitly constrained. These results enrich our understanding of the role played by the PLM during sequence design and illustrate the need for further work in determining the utility of such methods.

## 2 Methods

### 2.1 Enabling Backpropagation Design in ESMFold

Protein design by backpropagating through a structure prediction network has previously been reported for AlphaFold [9, 10, 11]. Our approach to enabling design of input sequences in the ESMFold network, which is built on OpenFold [15] and is outlined in Figure 1, is no different: design is achieved by minimizing the loss gradient with respect to the input rather than to the weights. Several changes in the source code were introduced to enable this functionality.

**Figure 1:**
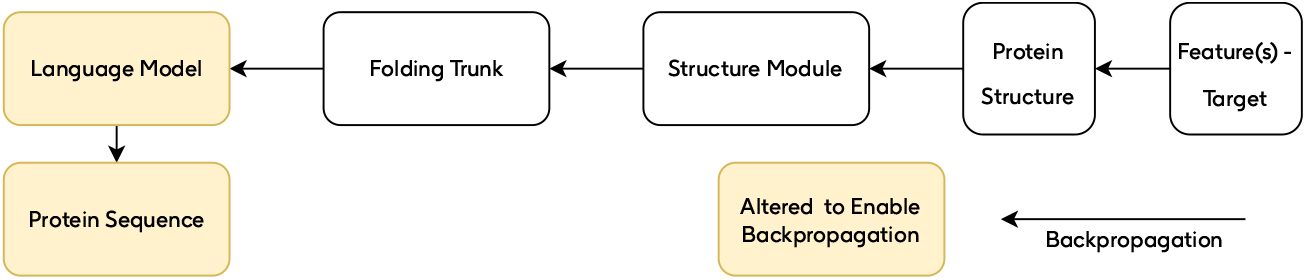
Schematic of dataflow through ESMFold. ESMFold infers structure from a protein sequence via two major elements: a protein language model (ESM2) and a structure module. In the language model, the string sequence is tokenized to integers. To enable backpropagation, the integers are converted to one-hot encoded floats and corresponding integer operations are converted to be compatible with the one-hot matrices rather than the integer vectors. Following this conversion, the computation graph can update amino acid identities based on a gradient calculated against any loss function.

Primarily, the input needs to be amenable to differentiation. In ESMFold, input sequences are tokenized as a vector of integers and prepended with a beginning of sequence (<BOS>) token. We converted the input from a vector of length *L* to a one-hot encoded matrix of shape (*N, L*), where *N* is the number of tokens and *L* is the sequence length plus the <BOS> token. Any downstream operation had to be converted to a corresponding differentiable operation on a matrix (*e*.*g*. embedding look up tables became matrix multiplications).

Next, a discrete representation of the amino acid sequence must be maintained. While the above technically enables backpropagation design, naively updating the inputs to minimize loss will result in continuous representations of discrete amino acids, which is not representative of reality. There are several approaches to steer the design towards reality that we investigated: (1) taking the arguments of the maxima (argmax) is non-differentiable but can be made differentiable by the “straight-through” trick [16], (2) annealing the Softmax function with a temperature is differentiable but only yields discrete solutions at *T* = 0, and (3) sampling the Gumbel-Softmax distribution with the “straight-through trick” which has been shown to perform better than the “straight-through” trick alone [17].

We find that annealing the Softmax function with a scaling temperature provides a sufficiently smooth gradient for convergence. Although we caution that this may depend on the loss function and optimizer.

The final components to backpropagation design are a loss function, an optimizer, and the algorithm itself, which will vary with the design task. We empirically evaluated the most appropriate options using two benchmark tasks of interest: backbone hallucination, where we optimized for maximum predicted local distance difference test (pLDDT, which measures confidence of the predicted structure), and backbone design/diversification, where we minimize root mean squared deviation (RMSD, which quantifies similarity to a reference structure) and maximize pLDDT. Based on the results of these tests, we elected to use Adam with a learning rat ≤ e 0.05. Stochastic gradient descent (SGD) did not converge as well as Adam, while AdamW performed about the same (Supplemental Figure 1). We found that raising the learning rate accelerated convergence when minimizing pLDDT but worsened performance when minimizing RMSD.

### 2.2 Improving Memory Limitations

One striking limitation of backpropagating through ESMFold versus AlphaFold is that a large difference in memory consumption is introduced by the presence of a 3 billion parameter language model in ESMFold. On a single A6000 GPU (48 GB), designing proteins greater than 100 residues is not possible. One potential solution to this is to split (shard) the model across multiple GPUs. In a naive test, splitting the ESM2 language model and ESMFold structure module each to their own GPU allowed the design of proteins up to 150 residues. Further sharding should permit increasing lengths, but users should be aware scaling is seemingly quadratic (Supplemental Figure 2). Alternatively, the design algorithm can be run on CPU but at roughly a 60-fold increase in runtime.

### 2.3 Comparison Runs in ColabDesign

We ran AFDesign with the [961c7af version of ColabDesign](https://github.com/sokrypton/ColabDesign) locally to make comparisons. Comparisons were run with the same number of steps as the ESM-Design strategy outlined above. The only notable difference was an additional “contact” term in the AFDesign loss function. Without this term, most AFDesigns runs tended to converge to extended alpha helices.

## 3 Results

### 3.1 Uniform Initialization Accelerates Design Convergence

In the simplest case, we sought to design for diverse, well-folded structures, maximizing pLDDT. We found that within 300 iterations, ESMFold hallucinated proteins with pLDDT values greater than 90, indicating very high confidence at the per-residue level. Although, it typically took as few as 100 optimization steps to converge to a unique fold. When we seeded a search against the PDB with a sample batch of 100-residue proteins designed for high pLDDT, we found no matches (FoldSeek version 5-53465f0 [18]). Several exemplars are shown in Figure 2. Broadly, this performance matched that of AFDesign.

**Figure 2:**
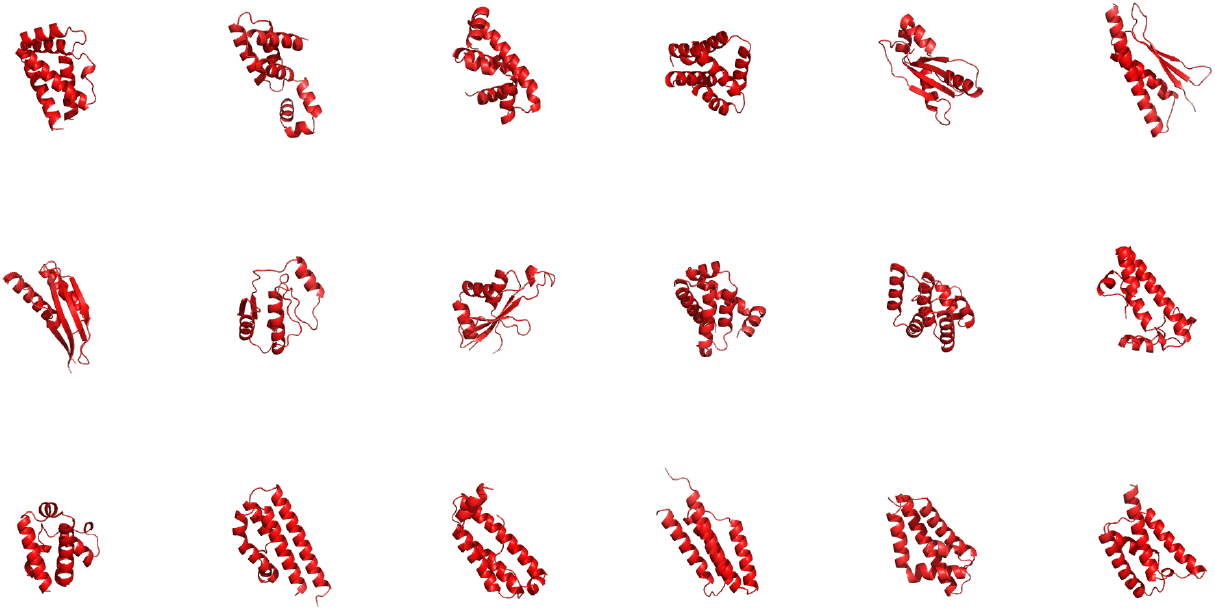
Diverse samples maximizing pLDDT, colored by pLDDT (blue - white - red). Structures are diverse although alpha is preferred over beta. All structures have pLDDT ∼ 90 across most positions.

During this initial trial, we observed that sequence initialization affected convergence, which has previously been reported [9]. To investigate more thoroughly we ran five initialization trials: (1) sampling the Gumbel distribution, as is default in AFDesign, (2) discretely sampling amino acids, each with probability 0.05, (3) discretely sampling amino acids, based on their fraction in UniRef90, (4) sampling a standard normal distribution, and (5) sampling a Gaussian distribution centered at 0.05 with a standard deviation of 0.02. The latter four sampling approaches were chosen to compare convergence when the model was given a distribution more or less similar to the one on which it was trained (UniRef90). The initialization trials were capped at 100 optimization steps to test early convergence.

Of these five approaches, we observed faster convergence when initializing with distributions more closely matching the training regime (*e*.*g*. discrete uniform or by sampling proportionally to occurrence rates in UniRef90). Whereas Gumbel initializations yielded a broad range of pLDDT values (from 40 to 60) after 100 iterations of ESM-Design, random initializations achieved pLDDT values between 70 and 90 (Figure 3). AFDesign fared slightly better, albeit with the caveat that uniform-initialized sequences typically converged to extended alpha helices, which is not practically useful. Gumbel-initialized sequences did not have this issue unless the “contact” term weight in the AFDesign loss function was set to zero. ESM-Design was also susceptible to this quirk, but only when initializing from the Gaussian distribution. Thus, we would recommend testing multiple initialization schemes at the start of each design project.

**Figure 3:**
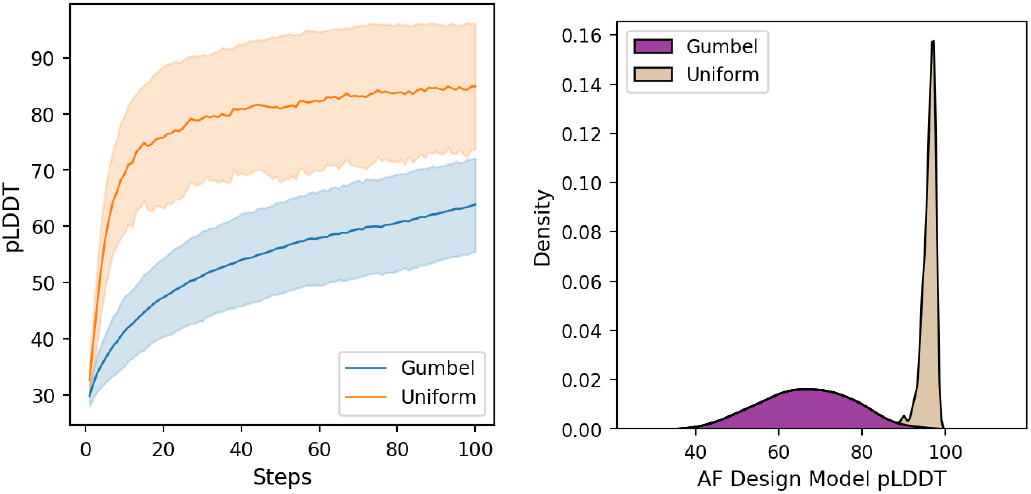
(Left) Average and standard deviation of pLDDT over the course of 100 design trajectories with ESM-Design initialized in one of two ways (discrete uniform sampling of all 20 amino acids or from the standard Gumbel distribution). (Right) The distribution of design pLDDTs following an abbreviated three stage design trajectory (50, 50, 10 iterations in each stage) with a 50 step “soft” warmup using AFDesign. Each distribution contains 100 independent design trajectories.

### 3.2 Design with RMSD Loss is More Challenging

Designing for high pLDDT is relatively trivial and serves primarily to provide novel backbones. A vastly more useful objective would be to design for specific structures or substructures. To that end, we implemented a differentiable Kabsch alignment algorithm [19], enabling an RMSD-based loss function. We note, however, that FAPE loss and distogram loss are also valid choices. We did not scale RMSD in the loss function, which caused RMSD loss to contribute 10-fold more than pLDDT; while perhaps suboptimal, this provides a starting point for fine-tuning the relative weights. We tested the loss function on a fibronectin domain (PDB: 1TEN), which has been reported to be challenging for MCMC approaches [20]. While convergence was slower than for pLDDT loss alone, designs eventually achieved alpha-carbon RMSD values below 2 Angstroms (Figure 4). It took five times more optimization steps to reach these RMSD values as it did to reach pLDDT values exceeding 90. However, we found that low RMSD values were not guaranteed, indicating that further investigation of the overall design strategy was necessary (*e*.*g*. designing in continuous space until RMSD reaches a certain threshold then switching to discrete space).

**Figure 4:**
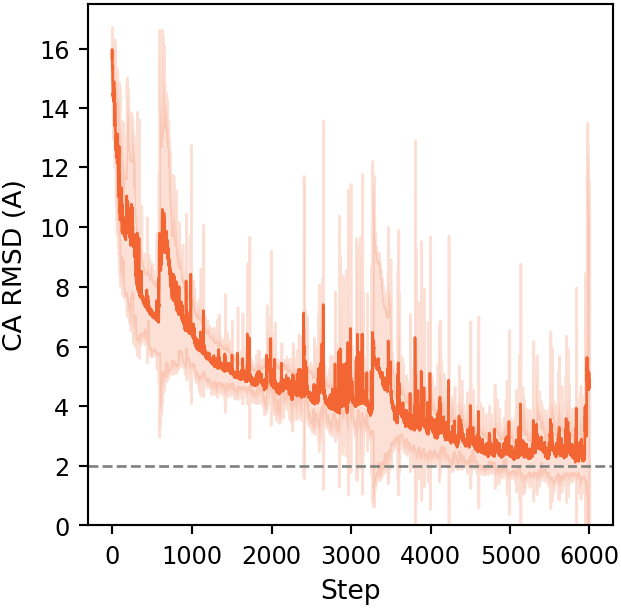
Sample traces of alpha-carbon RMSD with respect to 1TEN for five design trajectories. Bold line indicates the mean, while the outline shades one standard deviation.

Optimizing a sequence for a particular structure allows us to demonstrates how hallucinated sequences differ between ESMFold and AlphaFold. Specifically, we modified the sequence of the 76-residue protein ubiquitin to maximize pLDDT and minimize RMSD to its crystal structure (PDB: 1UBQ). Running ESM-Design for 100 independent trajectories of 300 steps each we generated designs that achieved a median RMSD of 0.08 Å and pLDDT of 0.917. Remarkably, we found that the most frequent amino acid at all 76 positions among these designs matched those of naturally occurring ubiquitins retrieved from UniRef90 using Jackhmmer (hmmer.org) (Figure 5). In contrast, sequences generated by AFDesign matched only at 24 of 76 positions, despite similarly low median RMSD values. Moreover, designs generated by ESMDesign achieved a median sequence identity of 39.6% with naturally occurring ubiquitins, illustrating how these reflect unique, *de novo* designed sequences. Simultaneously, looking at the sequence alignment of the designs as a whole indicates that the language model gravitates toward the family consensus sequence when designing for a target crystal structure.

**Figure 5:**
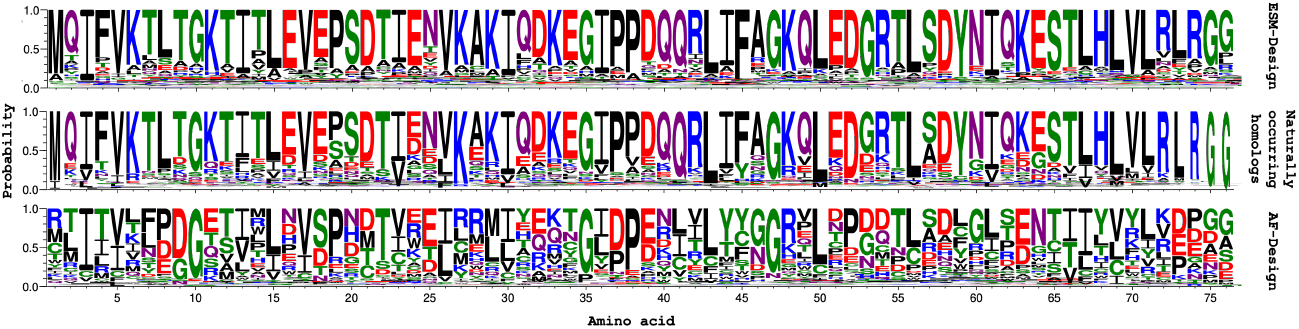
Sequence logo comparison of ubiquitin designs, plotting frequency rather than entropy. ESM-Design (top) more closely reproduces the UniRef90-derived MSA (middle) than AFDesign (bottom).

### 3.3 ESM-Design Diversifies Enzymes While Retaining Key Residues

Another practical application for hallucination is to produce slight variations on the sequence and structure of a protein active site. Diversification of an active site could, for example, enable catalysis with non-natural substrates. To that end, we tested the capacity for ESM-Design to generate minimal structural and sequence variants from a starting sequence that is known to fold and catalyze a particular reaction. When maximizing pLDDT on an undisclosed target enzyme approximately one-hundred and fifty residues in length, we found that 39.4% of design trajectories produced enzymes with key conserved residues in the active site while also diversifying the remainder of the protein sequence by 7-25%. Designs were structurally diverse as well, ranging in backbone alpha-carbon RMSD from 1.6 to 7.6 Angstroms (Figure 6), while conserving the active site geometry. Thus, ESM-Design provides a plausible alternative to minimal sequence and structural diversification methods, and respects conserved residues important to function without explicit user-defined constraints.

**Figure 6:**
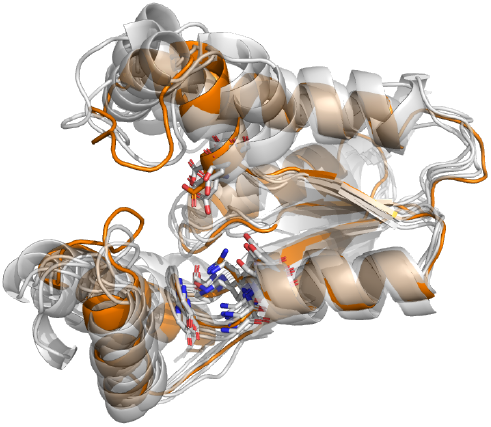
Alignment of ESM-Design-diversified models (white) to the target enzyme crystal structure (orange). The active site channel (center) is relatively unperturbed by the diversification while loops are more structurally variable.

### 3.4 ESM-Design Produces More Natural Designs

The original inspiration behind enabling design with the ESMFold neural network was to produce designs with more natural-like sequences, an observed deficiency of other backpropagation design strategies. We compared the amino acid usage rates of ESM-Design and AFDesign for a variety of initialization schemes, tasked with producing 100 well-folded (high pLDDT) sequences in 300 design steps. We discovered that initialization strongly affects amino acid usage rates for both ESM-Design and AFDesign (Supplemental Figure 3). Initializing by sampling from a UniRef-like distribution yields more native-like sequences from ESM-Design than AFDesign, possibly due to the coupled language model. AFDesign performed best, in terms of sequence similarity versus UniRef, when initialized by sampling from the Gumbel distribution.

Of course, we are not limited to design by backpropagation. The recently released structure-based sequence design method ProteinMPNN, a graph neural network trained on recapitulating sequences given a target backbone, has enabled rapid and reliable sequence design [12]. For each *de novo* designed protein backbone, we designed one sequence using ProteinMPNN version 1.0.1 with a temperature of 0.1 to test its effects on amino acid usage rates. Surprisingly, we found that MPNN produced less native-like sequences than either ESM-Design or AF Design (Figure 7 and Supplemental Figure 3). We speculate that this unexpected result is driven by the novel structures of the designs, which did not structurally align to anything in the PDB.

**Figure 7:**
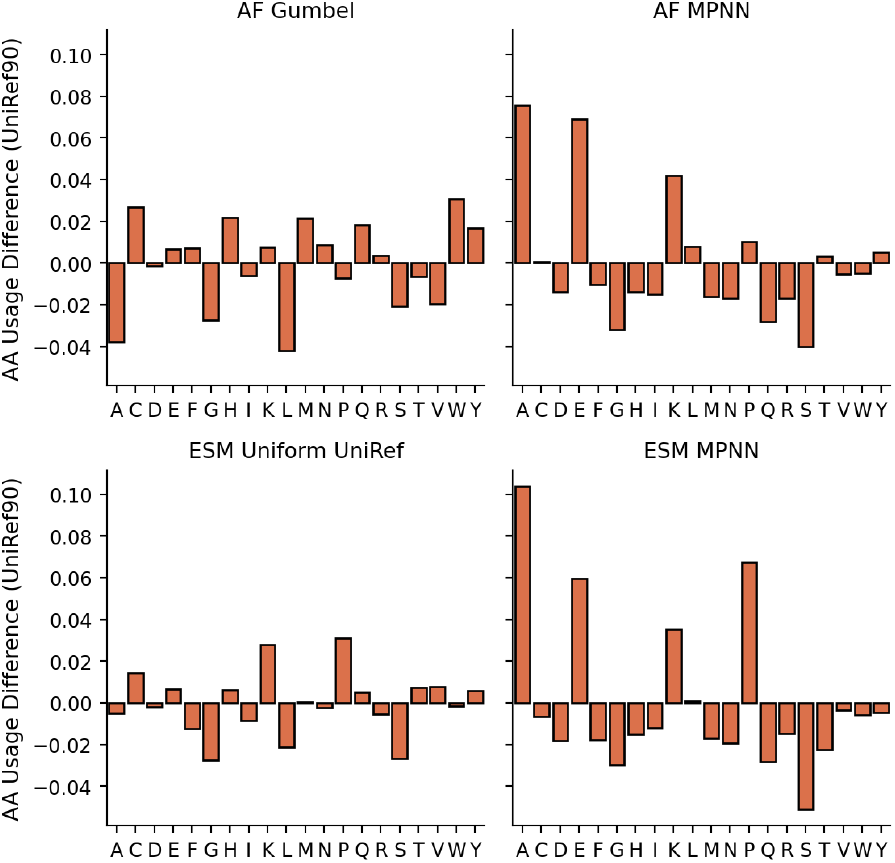
Differences in amino acid usage between design methods and UniRef90. Both design approaches prefer cysteine and proline more than is natural. MPNN shows preference for lysine and glutamate, and surprisingly alanine and proline, which is likely due to the atypical nature of the design structures.

To further assess whether the sequence profiles of these designs resembled those of naturally occurring proteins, we predicted *in silico* the likelihood of expression using our in-house model [21]. This model, which consists of a fine-tuned ProteinBERT [22] with a previously reported accuracy of 76%, provides an orthogonal assessment of whether sequences resemble those found in nature.

We found that ESM-Design sequences were twice as likely as AFDesign sequences to express when cutoffs were set to a reasonable threshold (Supplemental Figure 4). Interestingly, despite their unnatural amino acid usage rates, sequences designed with ProteinMPNN were predicted to express with greater confidence than those designed using either ESM-Design or AFDesign. This observation is consistent with previous reports of ProteinMPNN rescuing failed designs, and possibly explains the mismatch in amino acid usage between ProteinMPNN and UniRef. A second observation is that charged residues such as lysine and glutamate were overrepresented in the positive label class in the training set for the expression prediction model and appear to be preferentially introduced by ProteinMPNN, providing a plausible explanation for the increased probability of expression predictions. Thus, these results bolster the claim that the PLM component of ESM-Design improves the quality of designed sequences. They also support previous reports that ProteinMPNN rescued previously failed designs.

## 4 Discussion

This manuscript presents a brief overview of our attempt at enabling backpropagation design within ESMFold. This approach, which we term ESM-Design, is restricted by sequence length but offers several distinct advantages over alternatives such as AFDesign and ProteinMPNN. In designing with ESM-Design, model outputs were strongly affected by decisions around the use of discrete vs. continuous sequences, activation function, loss function, initialization conditions, optimizer, andoverall strategy. Moreover, complex interactions between these choices were not immediately clear. Nevertheless, several trends were apparent.

- Initialization affects the rate of convergence and ultimate amino acid usage. Unsurprisingly, initializing with discrete amino acids sampled with frequency akin to that in UniRef results in the fastest convergence and most natural-like designs. This contrasts with an inverted AlphaFold network, which instead benefited from a Gumbel distribution of amino acids during initialization.
- Several challenges became apparent when designing toward specific structures by minimizing RMSD. Designs converged more slowly, and unlike maximization of pLDDT, which routinely yielded high-confidence sequences, structures were not guaranteed to achieve RMSDs below a certain threshold after a fixed number of steps. Nevertheless, our results with ubiquitin suggest that including RMSD in the loss function constrained hallucination toward ubiquitin-like sequences. We note that although language models lacking a structural component have been used to achieve native-like sequences of specific families in the past, these required an additional fine-tuning step [7]. In contrast, our results suggest that no such fine-tuning would be required to hallucinate sequences for specific protein families, provided a reference structure is available.
- Catalytic amino acids at key positions were typically conserved during design without introducing explicit constraints, highlighting a role played by the PLM in navigating a complex fitness landscape defined by naturally occurring proteins with the same topology.
- Both ESM-Design and ProteinMPNN generated sequences predicted to express with high probability. AFDesign did not, consistent with previous reports [8, 12]. However, only ESM-Design generated sequences with amino acid frequencies matching those seen in UniRef.

In all four cases, we found that the PLM appeared to play a key role in regularizing sequence design away from unnatural compositions akin to those generated by an inverted AlphaFold network.

Nowhere was this more evident than when fine-tuning towards a pre-existing fold found in nature (Section 3.4). Conserved active site residues were typically unchanged, while surrounding residues were modified, indicating that the PLM has learned during training which residues are off-limits to design and which are not.

In conclusion, this work highlights how including a PLM can guide structure-based sequence design. While results are promising, they are necessarily limited to small proteins due to the huge memory footprint of the PLM and they are further muddled by the complex interplay between how sequences are initialized, and the choice of loss function and optimizer. Future research is needed to overcome memory limitations and identify an ideal training regime for design.

## Acknowledgments and Disclosure of Funding

This work would not have been possible without two fantastic open source projects. We thank Sergey Ovchinnikov for the permissive licensing and open development of ColabDesign. We thank Alex Rives and Meta for open sourcing ESM and ESMFold. We acknowledge the entire Protein Design and Informatics group at GSK for helpful discussions. Finally, we thank the GSK Fellows Program for research support.

## 5 Supplement

**Supplementary Figure 1:**
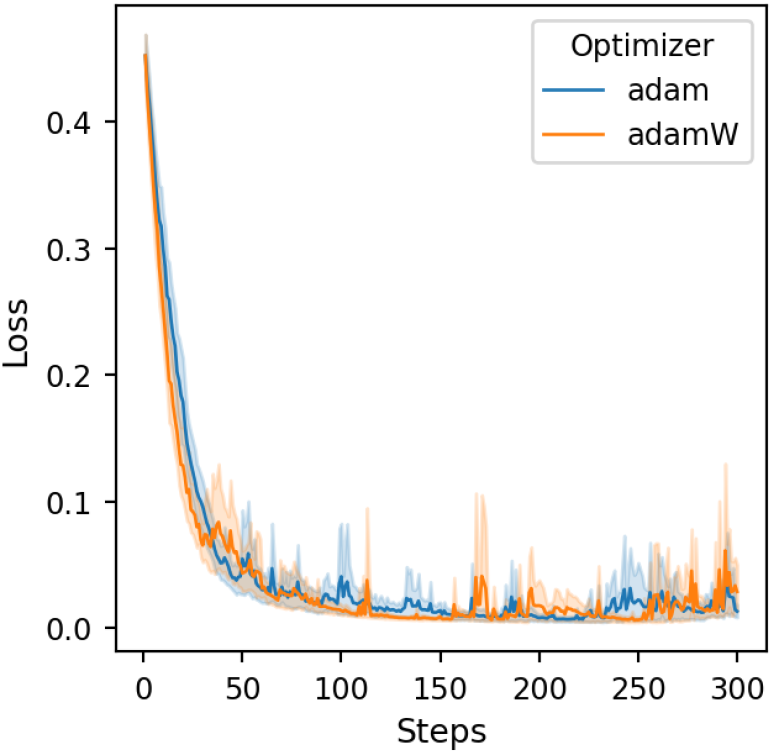
Comparison of pLDDT loss across ten runs with the Adam and AdamW optimizers in PyTorch. There is no clear difference between the two. SGD is not shown because it failed to converge in preliminary trials and was not further investigated.

**Supplementary Figure 2:**
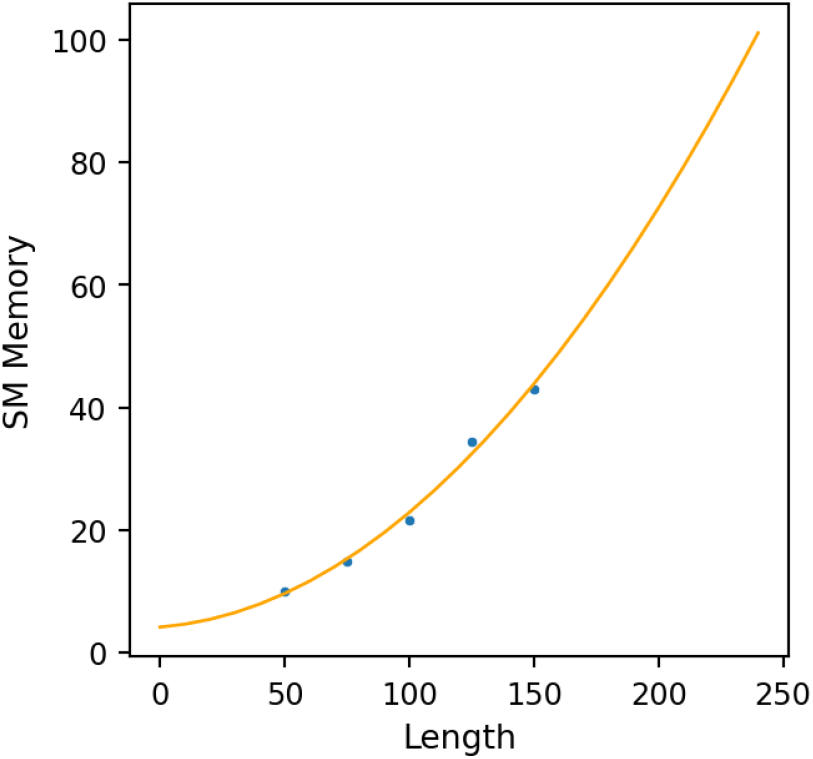
The structure module in ESMFold scales GPU RAM usage in *seemingly* quadratic fashion.

**Supplementary Figure 3:**
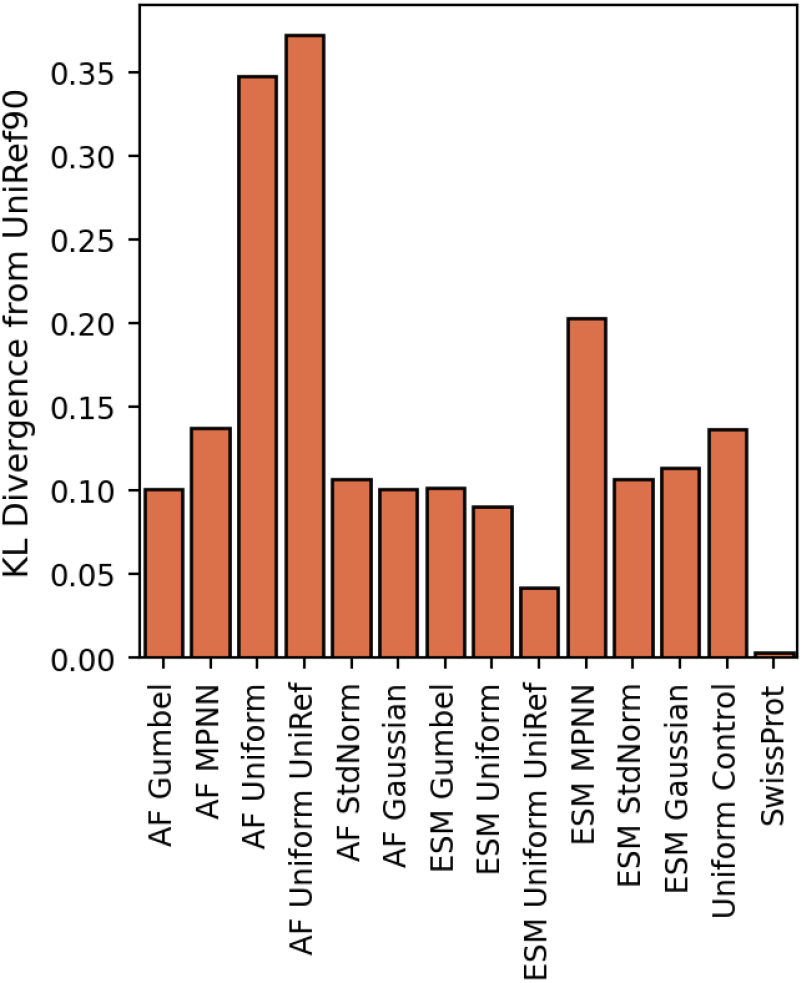
Using UniRef90 as a standard for “nature”, we compute KL divergence between the amino acid usage rates in UniRef90 and the designs produced by our various approaches. We find that ESM produces sequence distributions more similar to those observed in nature than other methods.

**Supplementary Figure 4:**
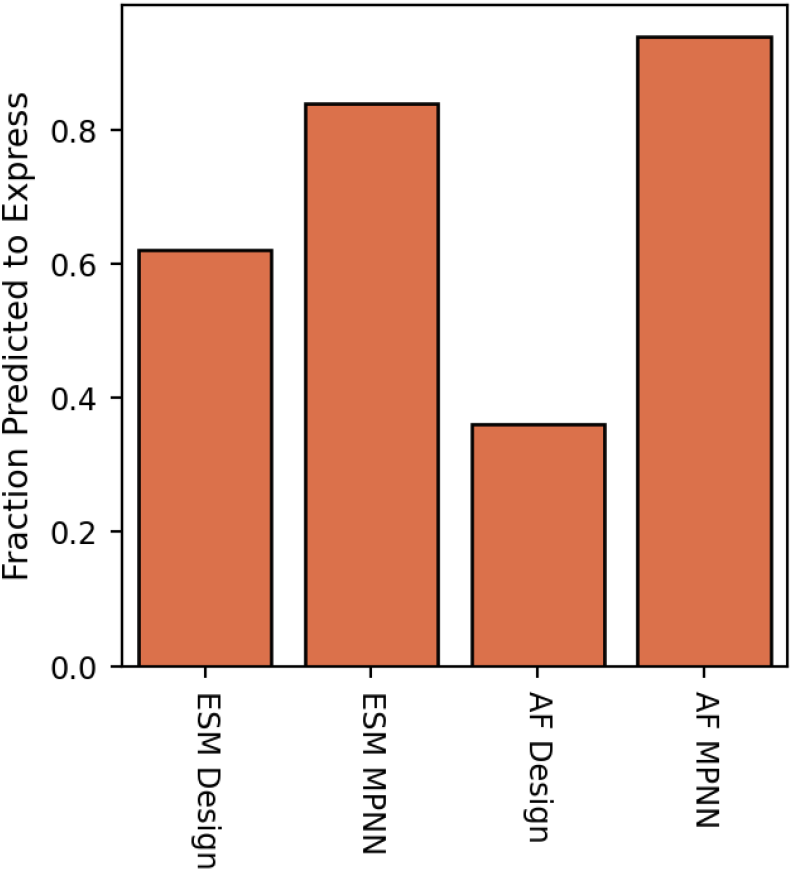
Fraction of sequences predicted to be expressing by an internal protein expression model.

## Notes

### Competing Interest Statement

The authors have declared no competing interest.

